# Linking stomatal function with photosynthetic light reactions and stress response in faba bean

**DOI:** 10.1101/2025.08.29.673109

**Authors:** Alexey Shapiguzov, Matleena Punkkinen, Tuomo Laine, Satu Engström, Pedro J. Aphalo, Hamid Khazaei

## Abstract

Faba bean (*Vicia faba* L.) is a key protein crop, but its cultivation and yield stability are hindered by a number of environmental stresses. Stomata regulate gas exchange between the plant and atmosphere, playing a central role in photosynthesis and mediating plant responses to a wide range of environmental stressors. This study aimed to investigate variations in leaf temperature as a proxy for stomatal function and photosynthetic regulations in faba bean, and to examine the response of this crop to short-term acute ozone (O_3_) exposure stress. Here, we used a high-throughput plant phenotyping (HTPP) platform to screen 196 faba bean genotypes for photosynthetic and stomatal function under controlled conditions. A subset of extreme genotypes, identified based on relative leaf tempreture from the initial screening, was exposed to a 450 ppb O_3_ treatment. Our results revealed strong positive relationship between photosynthetic efficiency and relative leaf temperature. A three-fold difference in relative leaf temperature was observed among genotypes. The O_3_ treatment caused signicantly less damage in genotypes with higher leaf temperature compared to those with lower leaf temperature (p<0.001). By combining a HTPP platform with elevated O_3_ stress treatment, we identified faba bean genotypes with contrasting stomatal responses to the O_3_ exposure. Our results advance understanding of the regulation mechanisms of photosynthetic light reactions and the role of stomatal function in modulating faba bean responses to environmental stressors.

## 1. Introduction

Faba bean is a globally important cool-season grain legume crop, valued for its high-protein, nutrient-rich seeds and its versatility as a source of food and animal feed (Duc, 1997; Khazaei and Vandenberg, 2020). In addition, its ability to fix atmospheric nitrogen reduces dependence on synthetic fertilizers, positioning it as a key species in low-input and environmentally sustainable cropping systems (Klippenstein et al., 2022). However, the cultivation of faba bean remains limited due to its susceptibility to a number of abiotic stresses such as water deficit (e.g., Khan et al., 2010; Muktadir et al., 2020), extreme temperatures (e.g., Bishop et al., 2016; Zhou et al., 2018), salinity (e.g., Bimurzayev et al., 2021; Tavakkoli et al., 2024), elevated tropospheric ozone (O_3_) levels (e.g., Otieno et al., 2022), and a wide range of biotic stresses (e.g., Rubiales and Khazaei, 2022). Increasing climate variability and environmental constraints emphasize the need to develop faba bean cultivars with improved resilience to adverse environmental factors (Mouritzen et al., 2025). Such improvements depend on a better understanding of morpho-physiological traits related to stress adaptation in this crop. Screening diverse faba bean germplasm collections for stress-relevant physiological traits accelerates the identification of pre-breeding resources for use in crop improvement programs aiming to ehance stress resilience (Adhikari et al., 2021) and provides critical insights into the mechanisms underlying stress adaptation in this species.

Stomata are central regulators of plant responses to many abiotic and biotic stresses, coordinating gas exchange, transpiration, and resistance to pathogens. Stomata control the exchange of water vapour and CO_2_ between leaves’ internal air spaces and the atmosphere. Thus, adjusting stomatal opening to the growing environment is crucial for plant growth and production. Reduced stomatal conductance can mitigate abiotic stress, but in most cases it concurrently decreases photosynthetic carbon assimilation by limiting CO_2_ flux into leaves (e.g., Salmon et al., 2020). Stomata are one of the main gateways for the entry of pathogens inside leaf tissues. They are thus intimately linked to plant immune reactions and biotic stress tolerance (Shapiguzov et al., 2012; Waszczak et al., 2018). Overall, under diverse environmental conditions responsiveness and control of stomatal aperture become critical for favourable carbon gain without excessive water loss as well as pathgen resistance. As water evaporation requires energy, leaves cool down with transpiration. As transpiration rate depends on stomatal conductance, differences in leaf temperature are correlated with differences in stomatal conductance. Leaf temperature has shown strong correlation with stomatal function in various plant species (e.g., Jones 1999; Chaerle et al., 2007; Sirault et al., 2009) including legumes species (Reynolds-Henne et al., 2010), and faba bean (Khan et al., 2010; Khazaei et al., 2013). Phenotypic variation in leaf temperature in a uniform environment may be a target for crop improvement efforts aiming to balance productivity and resilience.

Photosynthesis is tightly regulated at multiple levels, including adjustments in light harvesting and carbon metabolism. One key protective mechanism is non-photochemical quenching (NPQ), which allows plants to dissipate excessive light energy as heat under high light or stress conditions, preventing damage to photosynthetic apparatus. The magnitude and dynamics of NPQ, including its relaxation after transitions to lower light were shown to directly affect plant growth and production (Kromdijk et al., 2016; de Souza et al., 2022). The regulation of photosynthetic light reactions has not received sufficient attention in faba bean. More emphasis has been put on stomatal morphology and function and their role in stress response in faba bean (e.g., Darwish and Fahmy, 1997; Khazaei et al., 2013; Müllers et al., 2022), while the functional link between stomatal function and the regulation of photosynthesis has not been systematically characterised in this species.

Ozone (O_3_) serves as a model envronmental stressor that induces formation of reactive oxygen species (ROS), key signaling molecules involved in plant development and in responses to abiotic and biotic stresses (e.g., Kangasjärvi et al., 2005; Iyer et al., 2012; Krasensky et al., 2017). O_3_ gas enters leaf tissues though stomata, triggering immune reactions that under acute O_3_ exposures may cause cell death. O_3_ may be used to rapidly and uniformly impose stress for screening large numbers of genotypes for stomatal function (e.g., Waszczak et al., 2024). The phenotypic varation in response of stomatal function to acute O_3_ exposures in faba bean remain poorly studied.

In this study, we used high-throughput plant phenotyping (HTPP) tools to study the variation of leaf temperature and photosynthetic regulatory processes in 196 faba bean genotypes of different origins, using HTPP of diverse chlorophyll fluorescence parameters and leaf temperature. We also assessed the effect of acute O_3_ exposure on selected faba bean genotypes with contrasting leaf temperatures.

## 2. Materials and methods

### 2.1 Plant material

One hundred ninety-six faba bean genotypes were used in this study (Table S1). Seeds from all faba bean genotypes were selfed for at least one generation in an insect-proof greenhouse and then seeds used for this study. Seeds harvested from single plants were used.

### 2.2 Experiment 1: Screening faba bean germplasm

#### 2.2.1 Growing conditions

Experiment 1 was conducted in the climate-controlled glasshouse of the University of Helsinki, Viikki campus, Finland, using a randomized complete block design with four replicates in December 2023. Seeds of all genotypes were inoculated with *Rhizobium leguminosarum* biovar. *viciae* (faba bean strain, Elomestari Oy, Tornio, Finland) before sowing. Seeds were sown in black plastic pots (8 × 8 × 8 cm, apploximately 0.6 L) containing a 1:1 mixture of peat and vermiculite (Karkea Ruukutusseos, WR8014, Kekkilä Oy, Vantaa, Finland) containing all essential nutrients. Each replicate was placed on a seperate greehouse table. Soil moisture was maintained at field capacity with automatic table irrigation (three times per week) ensuring all plants were under well watered growing conditions. The photoperiod was set to 14 h of light and 10 h of darkness, with a day/night temperature regime of 21 °C/15 °C (±2 °C), and relative humidity was maintained at 60%. The main light source was high pressure sodium lamps (400W, Philips, Netherlands). Photosynthetic photon flux density (PPFD) at the canopy level was approximately 250 μmol m^−2^ s^−1^.

#### 2.2.2 HTPP phenotyping

Phenotyping based on multiple types of images was conducted on 4-week-old seedlings of 196 faba bean genotypes using PlantScreen SC Mobile System (PSI, Czech Republic) that was located in the darkened compartment of the same growth room, assuring minimal changes in temperature and humidily during the imaging process. Four individual seedlings per line were phenotyped. For each imaging session, twenty four randomly picked plants of different genotypes were arranged on a single tray, thermal and chlorophyll fluorescence (CF) images were acquired from a top view. Thermal images were captured immediately after loading the trays into the phenotyping chamber using the inbuilt InfraTec VarioCam thermal camera (1024 × 768 pixels). High-resolution grayscale images were analyzed in ImageJ, with relative leaf temperature expressed as grayscale intensity in defined leaf areas. To enable comparisons between trays, individual plant values were normalized to the total signal of all 24 plants corresponding to different randomized genotypes within the same tray.

After thermal imaging, plants were adapted to darkness for 30 min, then CF imaging was performed with the inbuilt 1.4-megapixel TOMI-2 camera (PSI). The imaging protocol included the measurments of Fo and Fm (dark-adapted minimal and maximal fluorescence, respectively), then a stepwise light ramp produced by the device’s LED array (cool white, 6500 K) and applied in 1.5-min steps of PPFD 200, 400, 600, 800, 1000, and 1200 μmol m^−2^ s^−1^, which was followed by a 7.5-min dark relaxation period. Saturating pulses (800 ms, ∼3000 μmol m^−2^ s^−1^ at canopy level) were triggered at the end of each light step to determine Photosystem II (PSII) quantum yields and NPQ. Maximum PSII efficiency (Fv/Fm) was calculated as (Fm − Fo)/Fm. NPQ was calculated as (Fm − Fm′)/Fm′ (Horton and Ruban, 1992), ETR as [(Fm’-Fs)/Fm’] * PAR * 0.5 * 0.84, where Fm’ is maximal flurescence and Fs is steady-state fluorescence for the given light intensity (Genty et al., 1989). NPQ relaxation rate was assessed by measuring partially relaxed NPQ after 1.5 min of darkness following the end of the light ramp. Data were processed using FluorCam 10 software (PSI).

### 2.3 Experiment 2: Acute ozone treatment

Based on results of the experiment 1 and seed stock avaiability, four genotypes from the lowest (#713, #138, #247, and #024) and three from the highest (#191, #020, and #088) relative leaf temperature groups were selected for the acute O_3_ exposure experiment.

#### 2.3.1 Growing conditions and ozone exposure

The plants were grown in the same plastic pots and medium as above, in the Aralab chambers (FITOCLIMA BIO, S600/D1200) under white LED (LED TUBE T5, 4000 K, Ledvance) light at 16 h of light (7:00-23:00) and 8 h of darkness photoperiod, with a day/night temperature regime of 21 °C/19 °C (±1 °C), day/night relative humidity 70 / 60%, and PPFD at the canopy level. The seedlings were exposed to elevated O_3_ treaments 18-21 days after sowing. For this, they were moved to ozonation chamber with the same day length, light intensity, temperature and relative humidity on the night before the treatment and in the following morning, after 2 h of light (7:00-9:00), exposed to O_3_ gas (450 ppb) for six hours (9:00-15:00). The experiment was performed twice with four replicates.

#### 2.3.2 Thermal imaging

The non-stress seedlings described in Section 2.3.1 were imaged with a thermal camera (382 x 288 pixels, 0.04 °C temperature resolution, PI-450, Optris, Germany) just before O_3_ stress experiments.

#### 2.3.3 Quantification of ozone-induced damage

After O_3_ exposure, all developed leaves were removed. For quantification of O_3_-induced damage in visible spectrum, the leaves were scanned with a flatbed scanner (Epson Perfection V750 Pro). The color balance of the scanned images was first changed to make the damaged areas more visible: cyan-red 30% towards red and magenta-green 30% towards magenta. Sizes of the discolored areas were then measured and compared to total leaf sizes with Fiji distribution package of ImageJ 2 (Rueden et al., 2017; Schindelin et al., 2012). The areas counted as undamaged had three main color profiles: light green (approximate color values r 61-65, g 56-60, b 6-9), mid-green (approximate color values r 44-48, g 44-48, b 4-7), and dark green (approximate color values r 25-29, g 41-45, b 9-12), while the areas counted as damaged had two main color profiles: light brown (approximate color values r 56-60, g 40-44, b 10-13) and red (approximate color values r 46-50, g 19-23, b 16-19). Alternatively, the cut leaves were stored for three days in cold conditions (+4 °C) moisturized with wet paper towels prior to scanning to further visualize the damage. No color correction was necessary for images of cold-stored leaves, and the color profiles were defined as: undamaged light green (approximate color values r 52-56, g 62-66, b 26-30) and green (approximate color values r 32-36, g 47-53, b 5-10), damaged grey (approximate color values r 34-38, g 38-41, b 15-17), and black (approximate color values r 10-14, g 13-17, b 2-5). For detection of O_3_-induced damage in near-infrared (NIR) spectrum, the detached leaves were imaged using the NIR system of the IMAGING-PAM M-Series (Walz) that employes reflection and backscattering of the 780 nm light beam. For false-colour imaging of Fv/Fm following the O_3_ exposure, the detached leaves were placed on moist background and acclimated to darkness for 10 min, after which Fv/Fm was determined with the IMAGING-PAM M-Series (Walz) using Walz measurment routines.

#### 2.3.4 Stomatal morphology

Stomatal density and size were measured on the middle part of the adaxial and abaxial surfaces of fully-expanded leaflets from non-stressed seedlings using the impression method. The stomatal imprints were taken from the same plants as were used for thermal imaging. Impressions were taken with Xantopren^®^ and its activator (Heraeus Kulzer GmbH, Germany), after which a thin layer of clear nail polish was applied to the impressions to create replicas. These replicas were then used for microscopic observations (Leica DMLB microscope with an attached ICC50W camera, Ernst Leitz Wetzlar GmbH, Heerbrugg, Switzerland). The number of stomata was calculated from at least three microscopic fields at 1218 × 914 µm (250x magnification), and stomatal size was measured at 500× magnification and converted to μm on eight stomata on the same leaflet samples as were used for stomatal density measurments.

### 2.4 Statistical analysis

Even though the layout in the greenhouse experiment followed a complete blocks design, the plants were randomized ignoring blocks during HTPP, thus a simpler one-way ANOVA model was used for data analysis. Principal component analysis (PCA) was done using genotype means. Figures were created in R version 4.5.1 (R Core Team, 2025) with packages ggplot2 (Wickham, 2016), ggpmisc (Aphalo, 2025) and segmented (Muggeo, 2025). Segmented-linear-regression models with one change point and the slope of the left side segment constrained to zero were fitted using genotype means as shown in Figure 3.

## 3 Results

### 3.1 Experiment 1

#### 3.1.1 Variation in relative leaf temperature and photosynthetic performance in faba bean germplasm

Highly significant phenotypic variation (p<0.001) in relative leaf temperature was observed among the studied 196 faba bean genotypes (Figure 1 and Table S2). A three-fold difference in relative leaf temperature was observed between genotype #713, which had the lowest relative leaf temperature, and genotype #191, which had the highest relative leaf tempreture.

**Figure 1.**
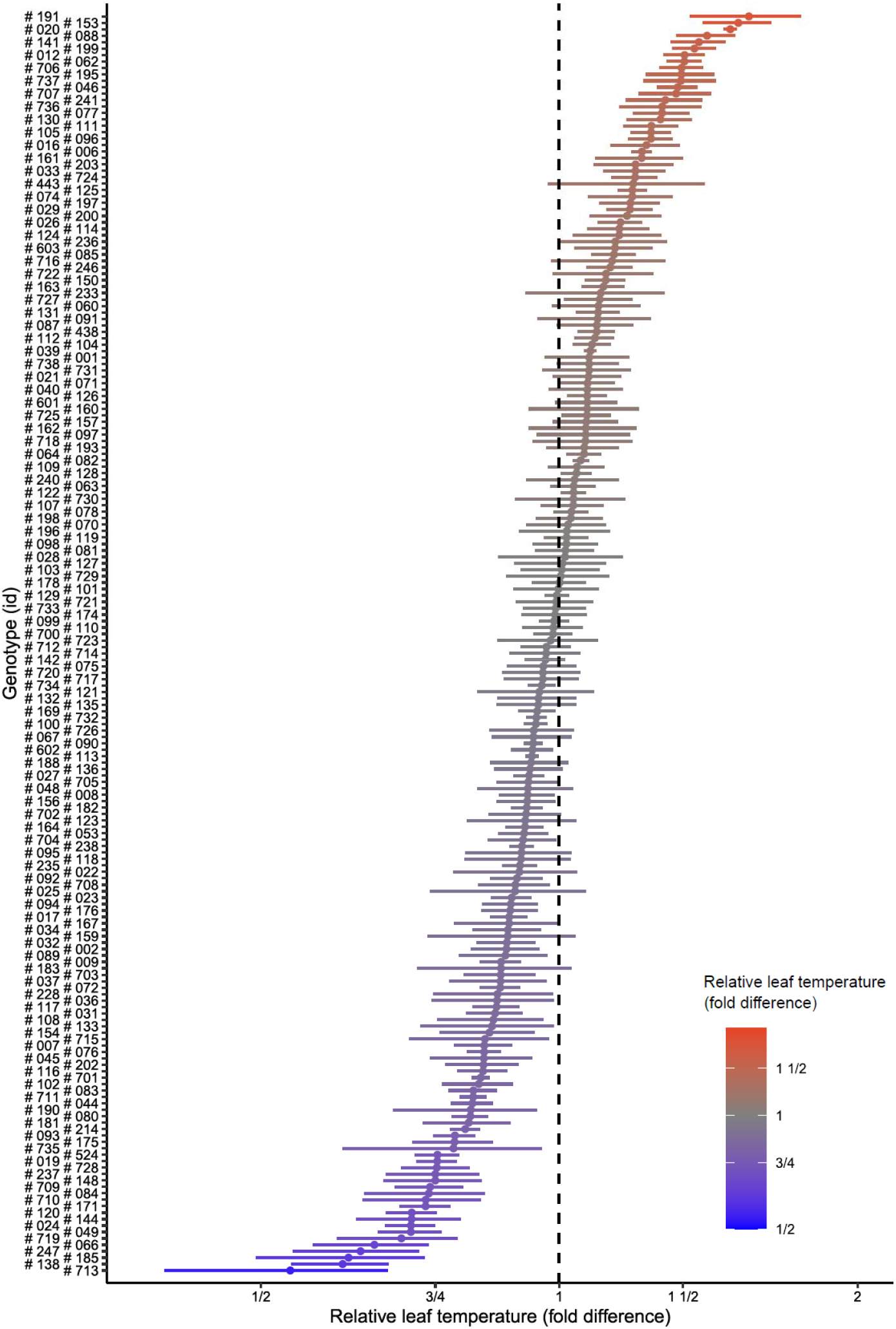
Relative leaf temperature in 196 faba bean genotypes. The vertical line indicates the mean relative leaf temperature. Colors indicate relative leaf temperature, ranging from highest (red) to lowest (blue).

The measurements of photosynthetic light reactions in the 196 faba bean genotypes revealed large variation in Photosystem II (PSII) quantum yields (QY), electron transfer rates (ETR) thorugh PSII and NPQ levels at different light intensities. These traits and the raw CF parameters used to calculate them are presented in Table S2. The NPQ and ETR across different tested light intensities are shown in Figure 2.

**Figure 2.**
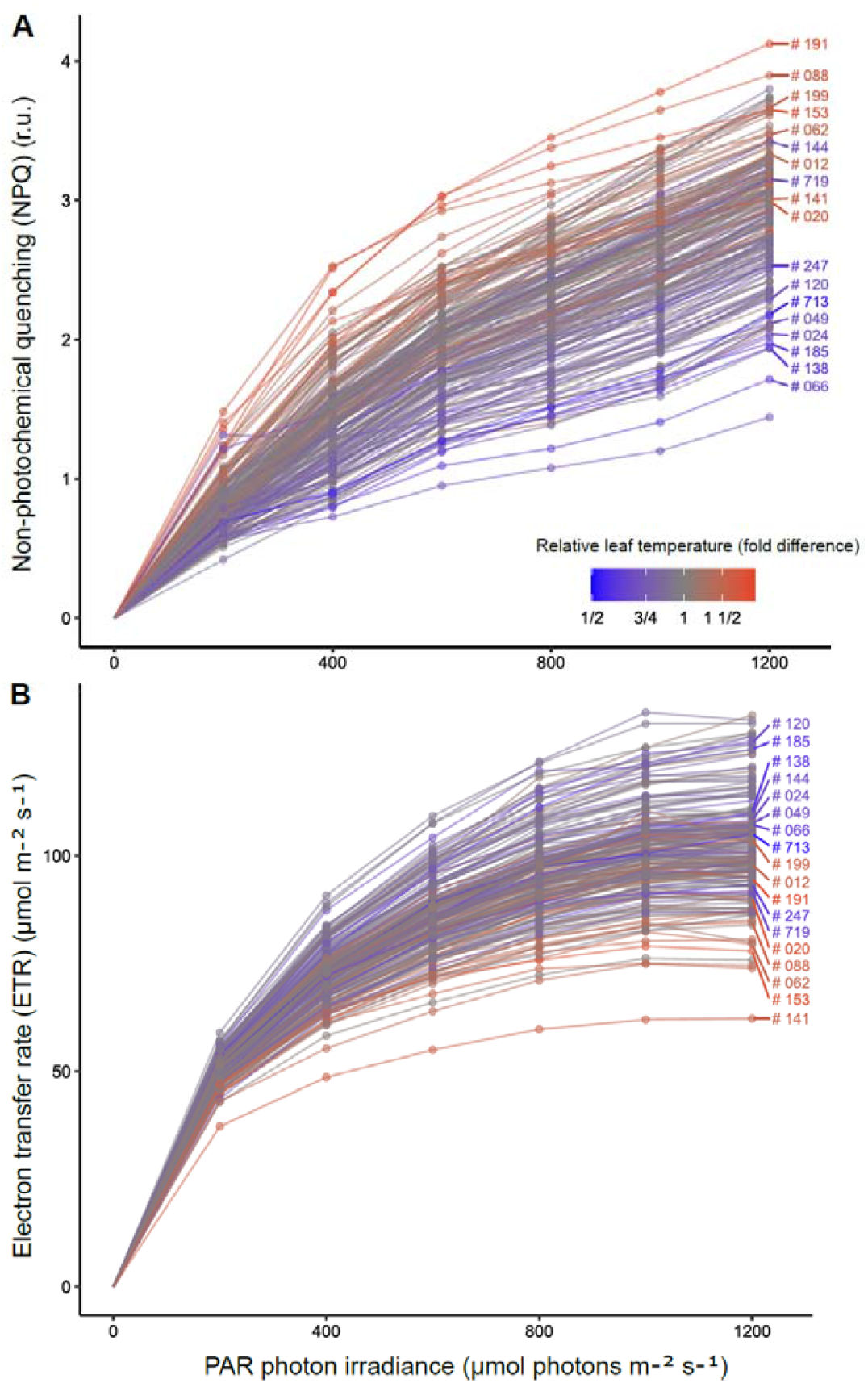
Variation of photosynthetic traits across the tested genotypes. NPQ (A) and ETR (B) were calculated under different light intensities in the 196 genotypes. Relative leaf temperature of individual genotypes is shown with the same color-coding as in Figure 1. Colors indicate relative leaf temperature, ranging from highest (red) to lowest (blue). The lines with the most extreme values of relative leaf temperature are labeled.

Substantial differences in the dynamics of NPQ and ETR between the genotypes with lower versus higher relative leaf temperature were detected (Figure 2). Across different light intensities, NPQ was lower in most of the coldest (low relative leaf temperature) genotypes and higher in the warmest (high relative leaf temperature) genotypes (Figure 2A). A reverse trend was observed in ETR, i.e., in the colder genotypes this parameter was overall higher than in the warm genotypes (Figure 2B).

#### 3.1.2 Relationships between relative leaf temperature and photosynthetic performance in faba bean germplasm

The above results suggested a link between the relative leaf temperature – a proxy for stomatal conductance, and photosynthetic light reactions that was likely related to limitation of photosynthesis by CO_2_ supply. We performed PCA on relative leaf temperature and 22 selected photosynthetic traits derived from CF imaging (Table S2) across the genotypes. The results are presented in Figure 3A. The two dimensions explained 79% of the total variance.

**Figure 3.**
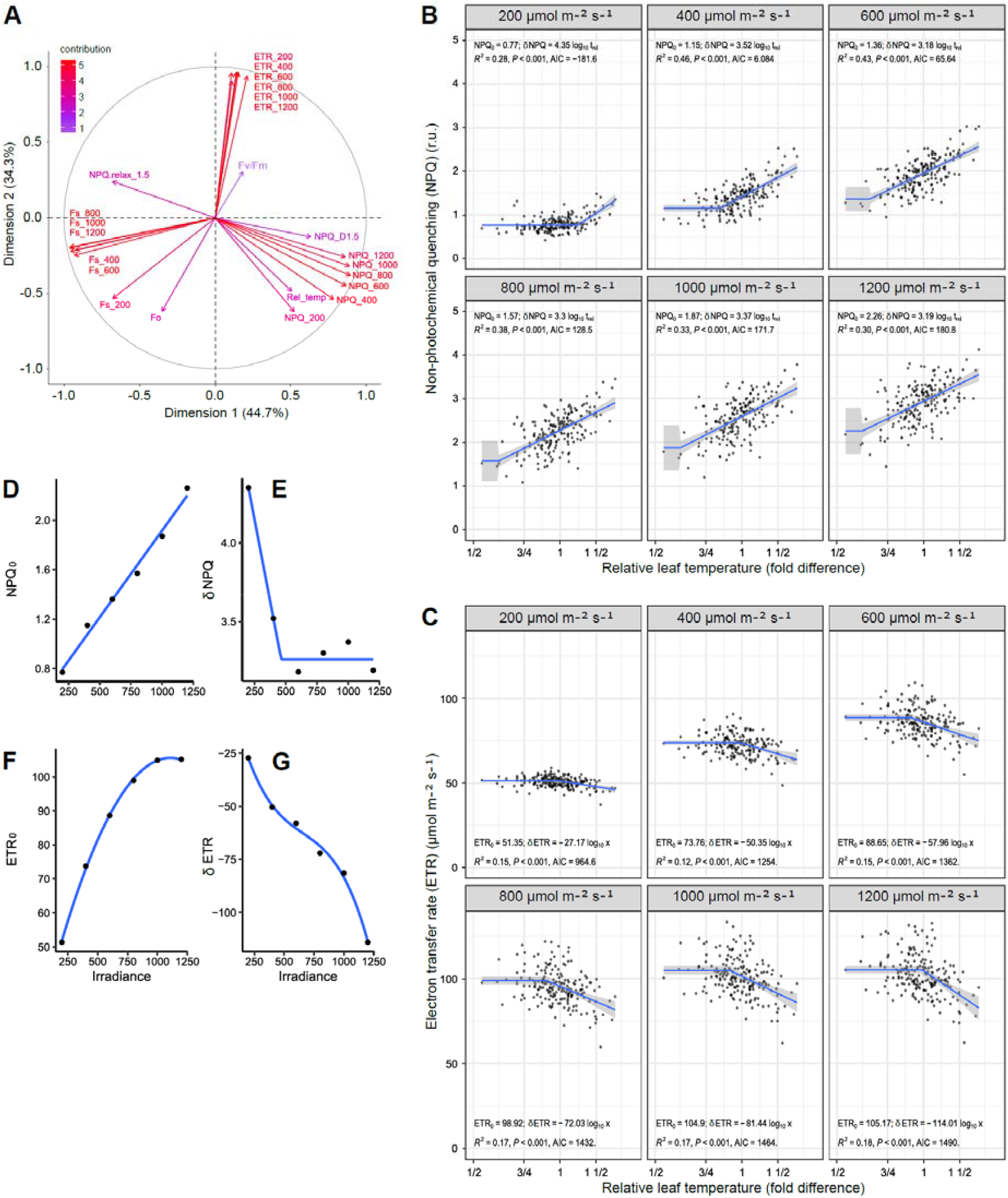
Relationship between photosynthetic functions and relative leaf temperature across 196 faba bean genotypes. (A) PCA of 23 phenotypic traits including relative leaf temperature, as well as QY, ETR and NPQ measured under different light levels. NPQ_D1.5 is the value of NPQ following 1.5 min of dark relaxation after the light ramp; NPQ_realx_1.5 is the ratio of NPQ_D1.5 over NPQ at 1200 µmol m^-2^ s^-1^; Fo is minimal and Fm is maximal dark-adapted CF; Fs is steady-state light-adapted CF; (*see* Table S2). (B) Correlations of relative leaf temperature with NPQ at different light intensities. (C) Correlations of relative leaf temperature with ETR at different light intensities. NPQ_0_ and ETR_0_ are the baseline values in the lines with low relative leaf temperatures, δNPQ and δETR are the changes in NPQ and ETR according to the changing log_10_ (relative leaf temperature). (D), (E), (F), and (G) Parameters estimates from (B) and (C), plotted vs. photon irradiance.

The vector of relative leaf temperature closely aligned with NPQ vectors at lower light intensities (200, 400, and 600 µmol m^-2^ s^-1^), suggesting a strong positive correlation that decreased under higher light intensities. This indicates that the genotypes with the higher relative temperature exhibited higher NPQ. In the model fits, variation in relative leaf temperature accounted for nearly 50% of the genotypic variation in NPQ (Figure 3B). As expected, NPQ increased with increasing irradiance (Figure 3B). Linear R^2^ between NPQ and leaf temperature increased with increasing irradiance until 400 and 600 µmol m^-2^ s^-1^ and then slightly decreased. This maximum corresponds to light intensities 2-3 times stronger than those during growth, suggesting that at these light intensities stomatal limitations on photosynthetic light reactions were strongest or more consistent among genotypes (Figure 3B).

In contrast to NPQ, the PCA vectors corresponding to ETR at all light intensities and the commonly used parameter Fv/Fm (the maximal yield of PSII photochemistry in the dark-adapted state) negatively correlated with the relative leaf temperature, indicating that higher leaf temperatures were associated with lower photosynthetic efficiency across all light intensities and in dark-adapted state (Figure 3C). However, relative leaf temperature was able to explain only between 10 and 16% of the variation in ETR among genotypes (Figure 3C). The partially relaxed NPQ (NPQ_relax_1.5) showed the same trend (Figure 3A). This indicated that higher relative leaf temperature may be associated with slower NPQ relaxation, which may reflect metabolic alterations imposed by more closed stomata. Baseline dark-adapted fluorescence level Fo also negatively correlated with leaf temperature, suggesting that the latter affected PSII function.

Overall, relative leaf temperature positively associated with NPQ, but weakly and negatively with actual photosynthetic performance (ETR and Fv/Fm) and NPQ relaxation rate. This indicated that under stomatal limitations to photosynthesis during brief exposure to high light faba bean seedlings activated photoprotection at the cost of photosynthetic efficiency. The change points separating the flat and sloped segments of the regression lines (Figure 3B and C) likely indicate a change in limiting factor. We hypothesize that these change points mark the onset of stomatal limitation to photosynthesis, i.e., the level of stomatal closure beyond which photosynthetic NPQ or ETR can no longer be maintained at their baseline values. Notably, with increasing light intensity, the change points shift along the x-axis towards lower relative leaf temperatures, suggesting that stomatal constraints on photosynthesis become more pronounced under higher PAR (Figure 3B and C). These light-dependent shifts differed between NPQ and ETR, indicating distinct mechanisms underlying their dependence on PAR and relative leaf temperature (Figure 3D-G).

### 3.2 Experiment 2

#### 3.2.1 Leaf temperature is linked to ozone stress response

To test whether the differences in relative leaf temperature, assumed to mainly reflect differences in stomatal conductance, correlated with altered tolerance to O_3_, we performed O_3_ gas fumigation in selected warm and cold faba genotypes form the experiment 1. Thermal imaging of the non-stressed plants confirmed the expected differences in leaf temperature among the selected extreme genotypes #713 and #191 (approximately 1 °C), although some genotypes identified in experiment 1 as cold (#247 and #024) did not follow the expected leaf temperature trend (Figure S1). Acute O_3_ exposure led to development of lesions in the colder, but not in the warmer genotypes identified in experiment 1. O_3_ lesions comprised large fraction of leaf area in colder genotypes (Figure 4A-C), while there was significantly less damage in genotypes with high relative leaf temperature (Figure 4A and B). No damage was detected in control plants that were not exposed to O_3_ (Figure S2). Stomatal density and size differed significantly among the genotypes (Figure S3A). The coolest genotype, #713, had significantly lower abaxial stomatal density than the other genotypes, however, adaixal stomatal density was not consistently different among genotypes. Genotype #713 also had the most open stomata based on stomatal pore width (Figure S3B).

**Figure 4.**
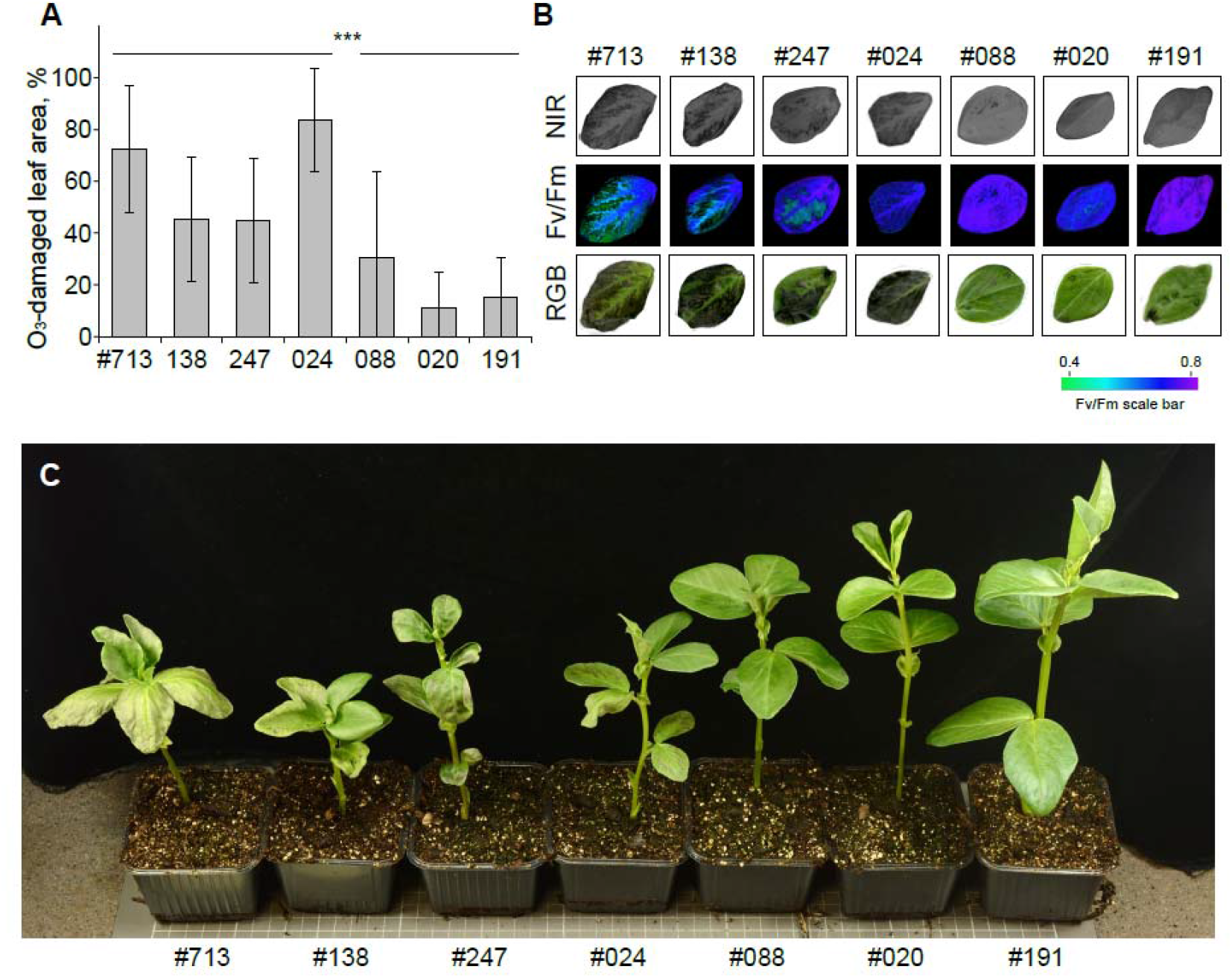
Tolerance to acute O_3_ exposure coincides with higher relative leaf temperature. (A) O_3_ lesions comprised large fraction of leaf area in colder genotypes. *** indicates a significant difference (p<0.001) between the mean of genotypes #713, #138, #247, #024 and the mean of genotypes #088, #020, #191 using *t*-test. The quantification is based on pooled data from two whole-experiment replicates, with four plants per genotype in each, but ignoring genotypes for the *t*-test. (B) Near-infrared (NIR) scans and photosynthetic yield (Fv/Fm) measurements were made in O_3_-treated leaves. After three days of cold storage, the same leaves were scanned with an RGB scanner. Results for non-stressed leaves are presented in Figure S2. (C) A photo of representative seedlings after the O_3_ exposure.

## 4. Discussion

This study explored phenotypic diversity for stomatal and photosynthetic-parameters in a large faba bean germplasm collection. We also demonstrated the link between stomatal function and stress response by exposing faba bean genotypes with extreme relative leaf temperature to the acute O_3_ stress. The combination of these approaches allowed us to establish a meaningful correlation between leaf temperature, as a proxy for stomatal function, and stress response in this species. This link has been reported in model plant species such as *Arabidopsis thaliana* (e.g., Wasczak et al., 2024), but remians less explored in crops (Yang et al., 2025). To our knowlegde, this is the first systematic study of these correlations across stomata-related phenotypes, photosynthesis and acute O_3_ stress tolerance in faba bean.

Our results revealed a strong relationship between relative leaf temperature and photosynthetic light reactions. The genotypes with warmer leaves exhibited higher NPQ and lower photosynthetic ETR, while the opposite pattern was observed in genotypes with cooler leaves. The relationships of ETR and NPQ with relative leaf temperature showed change points possibly attribuitable to irradiance-dependent thresholds in tolerance or changes in limiting factors. These patterns are consistent with pronounced stomatal limitations to photosynthesis in faba bean under irradiances doubling or more those during growth, whereby restricted CO_2_ availability leads to downregulation of photochemistry and upregulation of protective energy dissipation. Such limitations likely constrain carbon assimilation under fluctuating environmental conditions and may affect growth and yield potential (e.g., Harrison et al., 2020). Interestingly, the strongest correlation was observed at PPFD 400 and 600 µmol m^-2^ s^-1^, which was approximately two times higher than the growth light to which the seedlings were acclimated. The observed strong carbon limitation of photosynthesis in faba bean makes it a promising model for studying the interactions between water balance and photosynthesis. In particular, the role of the light environment needs to be considered when examining water stress responses, and analyses of photosynthesis, including CF traits. In the short term, light directly drives photosynthetic electron transfer, regulates stomatal opening and is the source of energy for evapotranspiration. In a longer time frame light irradiance and spectrum influence plants’ acclimation and stress tolerance (Aphalo and Sadras, 2022). Thus, illumination conditions may prove instrumental in such studies and in genotype screening. As faba bean cultivation expands into regions with variable climate regimes, a whole-plant physiological perspective will be essential for guiding the future breeding and production strategies. The physiological diversity revealed in this study provides a foundation for tailoring variety selection to regional growing conditions.

To our knowledge, this is the first systematic study in faba bean to characterize NPQ dynamics under varying light intensities across a broad genetic panel. The combination of CF and thermal imaging - especially when implemented using HTPP platforms - offers a promising path forward for screening complex physiological traits. These tools were shown to significantly enhance selection efficiency by providing quantitative, non-invasive indicators of performance under various stress conditions in this specis (e.g., Pedruzzi et al., 2020; Castillejo et al., 2021; Chen et al., 2023; Yang et al., 2024; Poque et al., 2025). Our results establish a physiological framework for identifying and developing more efficient, stress-resilient faba bean cultivars under controlled growing conditions.

Our results point to a possible role of stomatal function in the reponse to short-term acute O_3_ exposure in faba bean. Genotype #713, which had the lowest leaf temperature, showed more damage after acute ozonation, whereas genotype #191, with the highest leaf temperature, showed less damage. At the same time, stomatal imprint results indicated that #713 have slightly more open stomata. Reactions of plants to O_3_ have been addressed in diverse plant species (Yang et al., 2025). The effects of O_3_ on plants depend on both O_3_ concentration and duration of exposure. Chronic low-level exposure typically reduces photosynthesis, inhibits growth, and accelerates senescence without visible tissue damage. In faba bean such exposures only led to moderate decrease in photosynthesis and respiration (Aben et al.,1990), and changes in transpiration (Turcsanyi et al., 2000). In contrast, acute exposure to higher dose of O_3_ as used in the present study can induce cell death and visible lesions in sensitive plants (Vainonen and Kangasjärvi, 2003). O_3_ enters plant tissues through stomata and generates ROS in the apoplast, thereby mimicking pathogen-induced immune response (Kangasjärvi et al., 2005; Iyer et al., 2012; Krasensky et al., 2017) and triggering immune reactions leading to cell death (Shapiguzov et al., 2012; Waszczak et al., 2018). The magnitude of response depends on the effective dose of O_3_ within leaves and thus on stomatal aperture. Accordinlgy, we observed higher sensitivity to O_3_ of the faba bean genotypes with lower relative leaf temperature. This suggests that genotypes with higher stomatal opening may be more susceptible to O_3_-triggered damage, and potentially, to certain pathogens. Several pathogens, including powdery or downy mildew and rust fungi, bacteria and nematodes (e.g., Wu and Liu, 2022; Meddya et al., 2023; Hou et al., 2024), use stomata as entry points into host tissue. Our results therefore suggest that O_3_ sensitivity could serve as a useful tool for screening large faba bean germplasm for stress resistance.

Our experiments were conducted under climate-controlled growth conditions and controlled O_3_ stress treatments. However, under field conditions, a number of environmental variables such as light intensity, temperature, relative humidity, and wind speed will greatly influence physiological traits, particularly gas exchange parameters and leaf/canopy temperature (*reviewed in* Miguel Costa et al., 2013; Xu and Li, 2022). Experiments should be carried out under field growing conditions to validate controlled climate results and to better understand the genotype by environment interactions for such traits, which can provide valuable insights for breeders. Moreover, the screening methods described in this work under grenhouse and crontrolled environment chambers are useful tools for pre-screening large germplasm collections under controlled conditions for physiological and genetic studies.

Genetic control of photosynthesis-related traits such as photosynthetic rate, stomatal conductance and Fv/Fm has been shown to have complex inheritance in faba bean (Khazaei et al., 2019). The HTPP method described in this study, based on the non-invasive measurement of multiple physiological traits, will pave the way for screening a large diversity of faba bean genotypes and mapping populations to identify genomic regions governing stomatal function and photosynthesis-related traits in this species. Similar studies have previously used low-throughput phenotyping approaches to investigate stomatal morphology and function (Khazaei et al., 2014; Mandour et al., 2023). Using the identified faba bean genotypes with constracting stomatal function from this study, we are developing mapping populations to investigate the genetic and physiological basis of stomatal traits and photosynthetic activity. This will accelerate a better understanding of the mechanisms underlying the role of stomata in plant responses to environmental stresses.

## 5. Conclusions

In this proof-of-concept study, we combined high-throughput phenotyping with uniform stress treatment to screen a large faba bean germplasm. The results revealed interactions between stomatal function and photosynthetic light reactions, and allowed to characterize faba bean genotypes for their sensitivity and tolerance to acute O_3_ exposure. The described genetic resources and phenotyping approaches provide foundation for future physiological and molecular studies in faba bean.

## Acknowledgment

We thank Leena Grönholm for her kind assistance during the greenhouse experiments. We are also grateful to Varvara Shapiguzova for her help with the quantitative analysis of leaf temperature. Additionally, we thank the Nordic Genetic Resource Center (NordGen) and Prof. Wolfgang Link (Georg-August-Universität Göttingen) for providing the faba bean seeds.

## Authors contribution

Investigation: MP, TL, SE; Formal analysis: AS, MP, PJA; Writing – original draft: AS and HK; Writing – review & editing: all authors; Funding acquisition and Resources: HK and AS; Supervision and Conceptualization: HK and AS.

## References

Aben JMM, Janssen-Jurkovičová M, Adema EH (1990) Effects of low-level ozone exposure under ambient conditions on photosynthesis and stomatal control of Vicia faba L. Plant, Cell & Environment 13:463–469. 10.1111/j.1365-3040.1990.tb01323.x

Adhikari KN, Khazaei H, Ghaouti L, Maalouf F, Vandenberg A, Link W, O’Sullivan DM (2021) Conventional and molecular breeding tools for accelerating genetic gain in faba bean (Vicia faba L.). Frontiers in Plant Science 12:744259. 10.3389/fpls.2021.744259

Aphalo PJ (2025) ggpmisc: Miscellaneous Extensions to ‘ggplot2’. R package version 0.6.2.9001. 10.32614/CRAN.package.ggpmisc

Aphalo PJ, Sadras VO (2022) Explaining preemptive acclimation by linking information to plant phenotype. Journal of Experimental Botany 73:5213–5234. 10.1093/jxb/erab537

Bimurzayev N, Sari H, Kurunc A, Doganay KH, Asmamaw M (2021) Effects of different salt sources and salinity levels on emergence and seedling growth of faba bean genotypes. Scientific Reports 11:18198. 10.1038/s41598-021-97810-6

Bishop J, Potts SG, Jones HE (2016) Susceptibility of faba bean (Vicia faba L.) to heat stress during floral development and anthesis. Journal of Agronomy and Crop Science 202:508–517. 10.1111/jac.12172

Castillejo MÁ, Villegas-Fernández ÁM, Hernández-Lao T, Rubiales D (2021) Photosystem II repair cycle in faba bean may play a role in its resistance to botrytis fabae infection. Agronomy 11:2247. 10.3390/agronomy11112247

Chaerle L, Leinonen I, Jones HG, Van Der Straeten D (2007) Monitoring and screening plant populations with combined thermal and chlorophyll fluorescence imaging. Journal of Experimental Botany 58:773–784. 10.1093/jxb/erl257

Chen X, Zhong N, Luo Y, Ni Y, Liu Z, Wu G, Zheng T, Dang Y, Chen H, Li W (2023) Effects of strontium on the morphological and photosynthetic physiological characteristics of Vicia faba seedlings. International Journal of Phytoremediation 25:811–821. 10.1080/15226514.2022.2110037

De Souza AP, Burgess SJ, Doran L, Hansen J, Manukyan L, Maryn N, Gotarkar D, Leonelli L, Niyogi KK, Long SP (2022) Soybean photosynthesis and crop yield are improved by accelerating recovery from photoprotection. Science 377:851–854. 10.1126/science.adc9831

Darwish D, Fahmy G (1997) Transpiration decline curves and stomatal characteristics of faba bean genotypes. Biologia Plantarum 39:243–249. 10.1023/A:1000301205458

Duc G (1997) Faba bean (Vicia faba L.). Field Crops Research 53:99–109. 10.1016/S0378-4290(97)00025-7

Genty B, Briantais JM, Baker NR (1989) The relationship between the quantum yield of photosynthetic electron transport and quenching of chlorophyll fluorescence. Biochimica et Biophysica Acta (BBA)-General Subjects 990:87–92. 10.1016/S0304-4165(89)80016-9

Harrison EL, Arce Cubas L, Gray JE, Hepworth C (2020) The influence of stomatal morphology and distribution on photosynthetic gas exchange. The Plant Journal 101:768–779. 10.1111/tpj.14560

Horton P, Ruban AV (1992) Regulation of Photosystem II. Photosynthesis Research 34:375–385. 10.1007/BF00029812

Hou S, Rodrigues O, Liu Z, Shan L, He P (2024) Small holes, big impact: Stomata in plant– pathogen–climate epic trifecta. Moleculat Palnt 17:26–49. 10.1016/j.molp.2023.11.011

Iyer NJ, Jia X, Sunkar R, Tang G, Mahalingam R (2012) microRNAs responsive to ozone-induced oxidative stress in Arabidopsis thaliana. Plant Signaling & Behavior 7:484–491. 10.4161/psb.19337

Jones H (1999) Use of thermography for quantitative studies of spatial and temporal variation of stomatal conductance over leaf surfaces. Plant, Cell & Environment 22:1043–1055. 10.1046/j.1365-3040.1999.00468.x

Kangasjärvi J, Jaspers P, Kollist H (2005) Signalling and cell death in ozone-exposed plants. Plant, Cell & Environment 28:1021–1036. 10.1111/j.1365-3040.2005.01325.x

Klippenstein SR, Khazaei H, Vandenberg A, Schoenau J (2022) Nitrogen and phosphorus uptake and nitrogen fixation estimation of faba bean (Vicia faba L.) in Western Canada. Agronomy Journal 114:811–824. 10.1002/agj2.20945

Khan HR, Paull JG, Siddique KHM, Stoddard FL (2010) Faba bean breeding for drought-affected environments: a physiological and agronomic perspective. Field Crops Research 115:279–286. 10.1016/j.fcr.2009.09.003

Khazaei H, O’Sullivan DM, Sillanpää MJ, Stoddard FL (2014) Use of synteny to identify candidate genes underlying QTL controlling stomatal traits in faba bean (Vicia faba L.). Theoretical and Applied Genetics 127:2371–2385. 10.1007/s00122-014-2383-y

Khazaei H, Street K, Bari A, Santanen A, Stoddard FL (2013) Do faba bean (Vicia faba L.) accessions from environments with contrasting seasonal moisture availabilities differ in stomatal characteristics and related traits? Genetic Resources and Crop Evolution 60:2343–2357. 10.1007/s10722-013-0002-4

Khazaei H, Wach D, Pecio A, Vandenberg A, Stoddard FL (2019) Genetic analysis of photosynthesis-related traits in faba bean (Vicia faba) for crop improvement. Plant Breeding 138:761–769. 10.1111/pbr.12716

Khazaei H, Vandenberg A (2020) Seed mineral composition and protein Content of faba beans (Vicia faba L.) with contrasting Tannin contents. Agronomy 10:511. 10.3390/agronomy10040511

Krasensky J, Carmody M, Sierla M, Kangasjärvi J (2017) Ozone and Reactive Oxygen Species. In eLS, John Wiley & Sons, Ltd (Ed.). 10.1002/9780470015902.a0001299.pub3

Kromdijk J, Głowacka K, Leonelli L, Gabilly ST, Iwai M, Niyogi KK, Long SP (2016) Improving photosynthesis and crop productivity by accelerating recovery from photoprotection. Science 354:857–861. 10.1126/science.aai8878

Mandour H, Khazaei H, Stoddard FL, Dodd IC (2023) Identifying physiological and genetic determinants of faba bean (Vicia faba) transpiration response to evaporative demand. Annals of Botany 131:533–544. 10.1093/aob/mcad006

Meddya S, Meshram S, Sarkar D, Rakesh S, Datta R, Singh S, Avinash G, Kumar Kondeti A, Savani AK, Thulasinathan T (2023) Plant stomata: An unrealized possibility in plant defense against invading pathogens and stress tolerance. Plants 12:3380. 10.3390/plants12193380

Miguel Costa J, Grant OM, Manuela Chaves M (2013) Thermography to explore plant– environment interactions. Journal of Experimental Botany 64:3937–3949. 10.1093/jxb/ert029

Mouritzen TW, Meurer KHE, Bornhofen E, Janss L, Weih M, Andersen SU (2025) Faba bean genetics and crop growth models – progress to date and opportunities for integration. Plant and Soil 10.1007/s11104-025-07459-7

Muggeo VMR (2025) segmented: Regression Models with Break-Points / Change-Points Estimation (with Possibly Random Effects). R package version 2.1-4. 10.32614/CRAN.package.segmented

Muktadir Md A, Adhikari KN, Merchant A, Belachew KY, Vandenberg A, Stoddard FL, Khazaei H (2020) Physiological and biochemical basis of faba bean breeding for drought adaptation. Agronomy 10:1345. 10.3390/agronomy10091345

Otieno M, Peters MK, Duque L, Steffan-Dewenter I (2022) Interactive effects of ozone and carbon dioxide on plant-pollinator interactions and yields in a legume crop. Environmental Advances 9:100285. 10.1016/j.envadv.2022.100285

Pedruzzi DP, Araujo LO, Falco WF, Machado G, Casagrande GA, Colbeck I, Lawson T, Oliveira SL, Caires ARL (2020) ZnO nanoparticles impact on the photosynthetic activity of Vicia faba: effect of particle size and concentration. NanoImpact 19:100246. 10.1016/j.impact.2020.100246

Poque S, Carlson-Nilsson U, Omer M, Himanen K, Khazaei H (2025) Exploring an automated indoor high-throughput phenotyping facility to investigate the response of faba bean to water stress. Research Square. https://www.researchsquare.com/article/rs-6461902/v1

R Core Team (2025) R: A Language and Environment for Statistical Computing. R Foundation for Statistical Computing, Vienna, Austria. 10.32614/R

Reynolds-Henne CE, Langenegger A, Mani J, Schenk N, Zumsteg A, Feller U (2010) Interactions between temperature, drought and stomatal opening in legumes. Environmental and Experimental Botany 68:37–43. 10.1016/j.envexpbot.2009.11.002

Rubiales D, Khazaei H (2022) Advances in disease and pest resistance in faba bean. Theoretical and Applied Genetics 135:3735–3756. 10.1007/s00122-021-04022-7

Rueden CT, Schindelin J, Hiner MC, DeZonia BE, Walter AE, Arena ET, Eliceiri KW (2017). ImageJ2: ImageJ for the next generation of scientific image data. BMC Bioinformatics 18:529. 10.1186/s12859-017-1934-z

Salmon Y, Lintunen A, Dayet A, Chan T, Dewar R, Vesala T, Hölttä T (2020) Leaf carbon and water status control stomatal and nonstomatal limitations of photosynthesis in trees. New Phytologist 226:690–703. 10.1111/nph.16436

Schindelin J, Arganda-Carreras I, Frise E, Kaynig V, Longair M, Pietzsch T, Preibisch S, Rueden C, Saalfeld S, Schmid B, Tinevez J-Y, White DJ, Hartenstein V, Eliceiri K, Tomancak P, Cardona A (2012) Fiji: an open-source platform for biological-image analysis. Nature Methods 9:676–682. 10.1038/nmeth.2019

Shapiguzov A, Vainonen JP, Wrzaczek M, Kangasjärvi J (2012) ROS-talk - how the apoplast, the chloroplast, and the nucleus get the message through. Frontiers in Plant Science 3:292. 10.3389/fpls.2012.00292

Sirault XRR, James RA, Furbank RT (2009) A new screening method for osmotic component of salinity tolerance in cereals using infrared thermography. Functional Plant Biology 36:970–977. 10.1071/FP09182

Tavakkoli E, Watts-Williams SJ, Rengasamy P, McDonald GK (2024) Eliciting the aboveground physiological regulation that underlies salinity tolerance in faba bean (Vicia faba L.). Environmental and Experimental Botany 226:105849. 10.1016/j.envexpbot.2024.105849

Turcsanyi E, Lyons T, Plochl M, Barnes J (2000) Does ascorbate in the mesophyll cell walls form the first line of defence against ozone? Testing the concept using broad bean (Vicia faba L.). Journal of Experimental Botany 51:901–910. 10.1093/jexbot/51.346.901

Vainonen JP, Kangasjärvi J (2015) Plant signalling in acute ozone exposure. Plant, Cell & Environment 38:240–252. 10.1111/pce.12273

Waszczak C, Carmody M, Kangasjärvi J (2018) Reactive oxygen species in plant signaling. Annual Review of Plant Biology 69:209–236. 10.1146/annurev-arplant-042817-040322

Waszczak C, Yarmolinsky D, Leal Gavarrón M, Vahisalu T, Sierla M, Zamora O, Carter R, Puukko T, Sipari N, Lamminmäki A, Durner J, Ernst D, Winkler JB, Paulin L, Auvinen P, Fleming AJ, Andersson MX, Kollist H, Kangasjärvi J (2024) Synthesis and import of GDP-L-fucose into the Golgi affect plant-water relations. New phytologist 241:747–763. 10.1111/nph.19378

Wickham H (2016) ggplot2: Elegant Graphics for Data Analysis. Springer-Verlag New York, USA. R package version 3.5.2. 10.32614/CRAN.package.ggpplot2

Wu J, Liu Y (2022) Stomata–pathogen interactions: over a century of research. Trends in Plant Science 27:964–967. 10.1016/j.tplants.2022.07.004

Xu R, Li C (2022) A review of high-throughput field phenotyping systems: Focusing on ground robots. Plant Phenomics 2022:9760269. 10.34133/2022/9760269

Yang W, Zhang Z, Yuan T, Li Y, Zhao Q, Dong Y (2024) Intercropping improves faba bean photosynthesis and reduces disease caused by Fusarium commune and cinnamic acid-induced stress. BMC Plant Biology 24:650. 10.1186/s12870-024-05326-8

Yang N, Cotrozzi L, Liu C, Qiao Q, Wang X, Zheng F, Hoshika Y, Nali C, Paoletti E, Pisuttu C, Risoli S, Pellegrini E (2025) Development trend, evolution of themes, gaps, and challenges of the research investigating ozone effects on vegetation. Environmental Reviews 33:1–13. 10.1139/er-2024-0038

Zhou R, Hyldgaard B, Yu X, Rosenqvist E, Magaña Ugarte R, Yu S, Wu Z, Ottosen C-O, Zhao T (2018) Phenotyping of faba beans (Vicia faba L.) under cold and heat stresses using chlorophyll fluorescence. Euphytica 214:68. 10.1007/s10681-018-2154-y

