## Supplementary Figures for "Linking stomatal function with photosynthetic light reactions and stress response in faba bean"

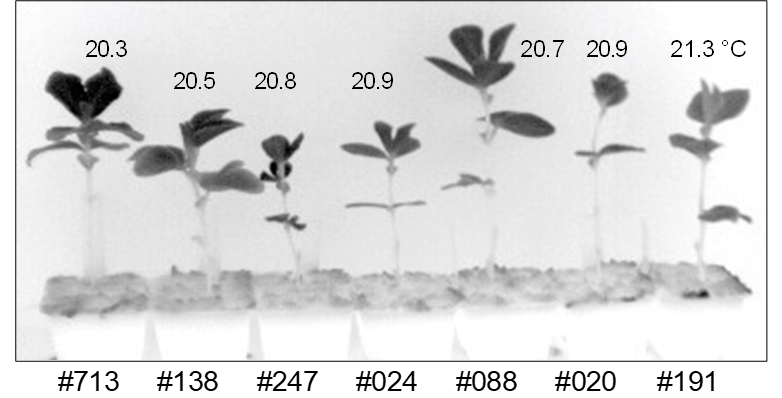

**Figure S1.** Thermal image of the non-stress seedlings (experiment 2).

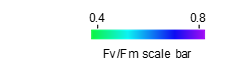

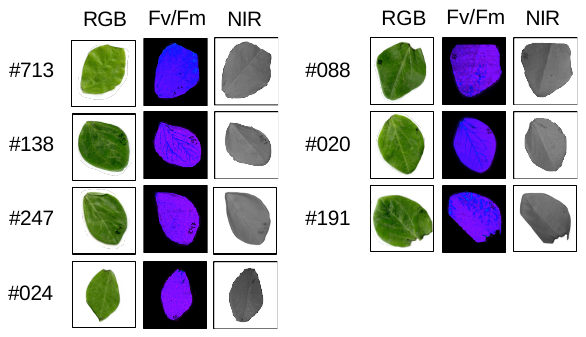

**Figure S2.** Excised control faba bean seedling leaves (experiment 2). Near-infra-red light scan (NIR) and photosynthetic yield (Fv/Fm) measurement were made in non-treated leaves at the same time as O_3_-treated leaves. Leaves were similarly scanned with a RGB scanner after three days of cold storage (RGB).

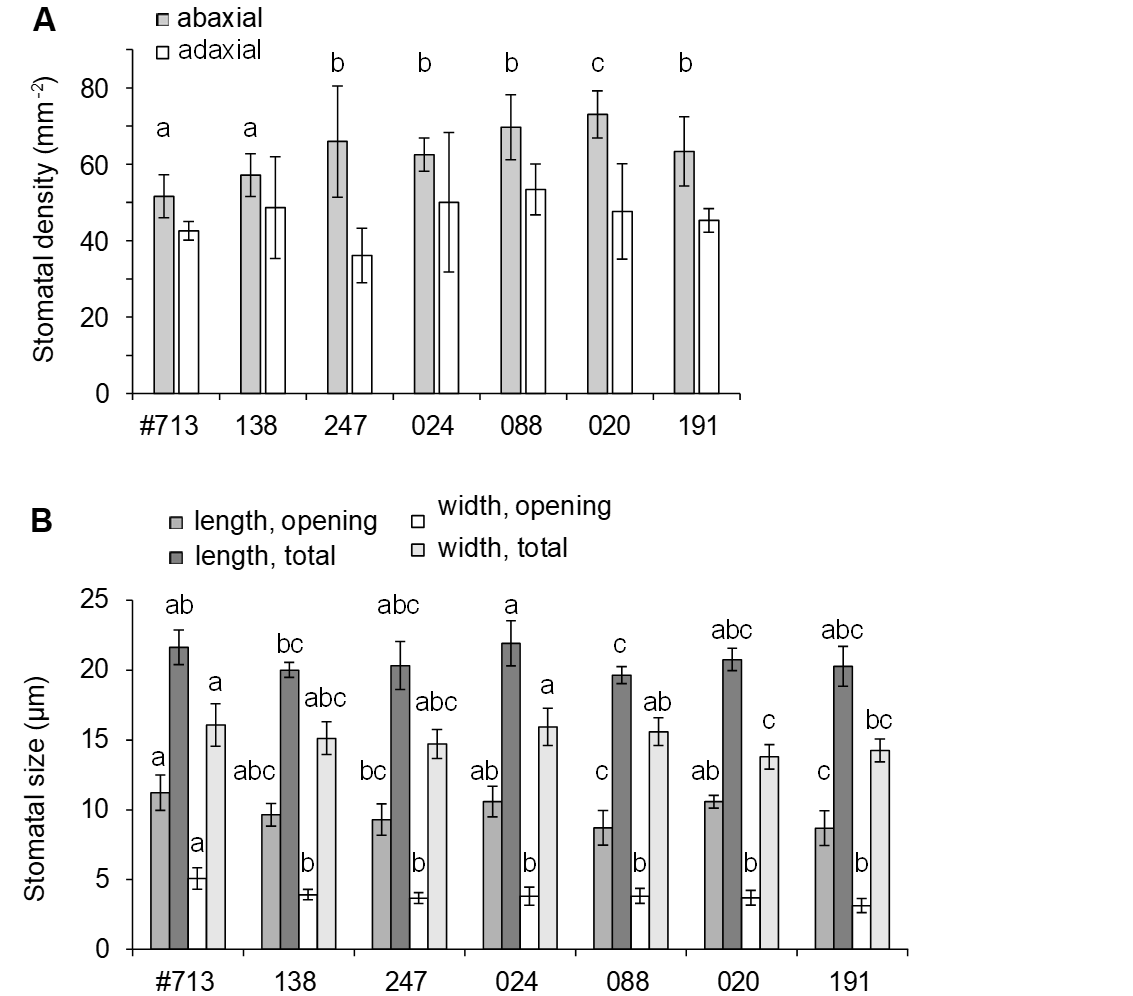

**Figure S3.** Stomatal density (A) and size (B) in faba bean seedling leaves (experiment 2). All data was acquired from surface imprints that were made from leaves attached to plants that were grown in growth chambers. Statistically significant groups (homogenous subsets determined with one-way ANOVA, post-hoc Tukey) for abaxial stomatal density and both dimensions of stomatal opening are marked with letters.
